# The conserved phosphatase GSP-2/PP1 promotes germline immortality via small RNA-mediated genome silencing during meiosis

**DOI:** 10.1101/273722

**Authors:** Katherine Kretovich Billmyre, Anna-lisa Doebley, Bree Heestand, Tony Belicard, Aya Sato-Carlton, Stephane Flibotte, Matt Simon, Megan Gnazzo, Ahna Skop, Donald Moerman, Peter Mark Carlton, Peter Sarkies, Shawn Ahmed

**Affiliations:** Department of Genetics, University of North Carolina, Chapel Hill, NC, 27599 USA; Department of Biology, University of North Carolina, Chapel Hill, NC, 27599 USA; Medical Research Council London Institute of Medical Sciences, Du Cane Road, London, W12 0NN, UK; Institute for Clinical Sciences, Imperial College London, Du Cane Road, London, W12 0NN, UK; Graduate school of Biostudies, Kyoto University, Kyoto, 606-8501, Japan; Department of Zoology, University of British Columbia, Vancouver, British Columbia, Canada; Department of Genetics, University of Wisconsin-Madison, Madison, WI 53706; Department of Opthamology, University of Washington, Seattle, WA, USA; Department of Biology, University of Rochester, Rochester, NY, USA

## Abstract

Genomic silencing can promote germ cell immortality, or transgenerational maintenance of the germ line, via mechanisms that may occur during mitosis or meiosis. Here we report that the *gsp-2* PP1/Glc7 phosphatase promotes germ cell immortality. We identified a separation-of-function allele of *C. elegans* GSP-2 that caused a meiosis-specific chromosome segregation defect and defects in transgenerational small RNA-induced genome silencing. GSP-2 is recruited to meiotic chromosomes by LAB-1, which also promoted germ cell immortality. Sterile *gsp-2* and *lab-1* mutant adults displayed germline degeneration, univalents and histone phosphorylation defects in oocytes, similar to small RNA genome silencing mutants. Epistasis and RNA analysis suggested that GSP-2 functions downstream of small RNAs. We conclude that a meiosis-specific function of GSP-2/LAB-1 ties small RNA-mediated silencing of the epigenome to germ cell immortality. Given that hemizygous genetic elements can drive transgenerational epigenomic silencing, and given that LAB-1 promotes pairing of homologous chromosomes and localizes to the interface between homologous chromosomes during pachytene, we suggest that discontinuities at this interface could promote nuclear silencing in a manner that depends on GSP-2.

**Author Summary:** The germ line of an organism is considered immortal in its capacity to give rise to an unlimited number of future generations. To protect the integrity of the germ line, mechanisms act to suppress the accumulation of transgenerational damage to the genome or epigenome. Loss of germ cell immortality can result from mutations that disrupt the small RNA-mediated silencing pathway that helps to protect the integrity of the epigenome. Here we report for the first time that the *C. elegans* protein phosphatase GSP-2 that promotes core chromosome biology functions during meiosis is also required for germ cell immortality. Specifically, we identified a partial loss of function allele of *gsp-2* that exhibits defects in meiotic chromosome segregation and is also dysfunctional for transgenerational small RNA-mediated genome silencing. Our results are consistent with a known role of *Drosophila* Protein Phosphatase 1 in heterochromatin silencing, and point to a meiotic phosphatase function that is relevant to germ cell immortality, conceivably related to its roles in chromosome pairing or sister chromatid cohesion.

## Introduction

Animals, including humans, are comprised of two broad cell types: somatic cells and germ cells. Somatic cells consist of many diverse differentiated cell types, while germ cells are specialized to produce the next generation of offspring. An important difference between these two cell types is that somatic cells undergo aging phenomena while the germ line is effectively immortal and capable of creating new “young” offspring [1]. Understanding the basis of immortality in germ cells may provide insight into why organisms age.

In *C. elegans*, disruption of pathways that promote germ cell immortality results in initially fertile animals that become sterile after reproduction for a number of generations. Many such *mortal germline* (*mrt*) mutant strains are temperature-sensitive, becoming sterile at 25°C but remaining fertile indefinitely at 20°C [2]. Mutations that cause a Mrt phenotype have been reported in two distinct pathways: telomerase-mediated telomere maintenance [3,4] and small RNA-mediated nuclear silencing [5–9]. Mutations in the PIWI Argonaute protein cause immediate sterility in many species.

However, disruption of the *C. elegans* Piwi orthologue PRG-1, which interacts with thousands of piRNAs to promote silencing of some genes and many transposons in germ cells, results in temperature-sensitive reductions in fertility and in an unconditional Mrt phenotype [6–12]. Multiple members of a nuclear RNA interference (RNAi) pathway that promotes maintenance of transgene silencing also promote germ cell immortality and likely function downstream of PRG-1/piRNAs [10,13]. One nuclear RNAi defective mutant, *nrde-2*, a number of heritable RNAi mutants, including *hrde-1*, and two RNAi defective mutants, *rsd-2* and *rsd-6*, only become sterile after growth for multiple generations at the restrictive temperature of 25°C [10,12–16]. These ‘small RNA-mediated genome silencing’ genes repress deleterious genomic elements via a small RNA-mediated memory of ‘self’ vs ‘non-self’ segments of the genome [13,17,18]. The transgenerational fertility defects of such mutants could reflect a progressive desilencing of heterochromatin due to changes in histone modifications downstream of small RNAs, which is modulated by histone modifications that occur in response to small RNAs, such as H3K4 demethylation and H3K9me2/3 [15,19].

Pioneering studies in *Neurospora* demonstrated that unsuccessful pairing of DNA during meiotic prophase can trigger small RNA-mediated genome silencing [20], and multigenerational silencing of hemizygous transgenes [21] suggests similar mechanisms may operate in *C. elegans*. Components of the *C. elegans* small RNA-mediated genome silencing machinery, such as the HRDE-1 and PRG-1 Argonaute proteins, are expressed throughout germ cell development [6,10,13,18]. RSD-2 displays a cytoplasmic localization in embryos but becomes a nuclear protein in adults [12], suggesting that aspects of small RNA-mediated genome silencing may be developmentally plastic.

Here we report the identification of a hypomorphic allele of *gsp-2*, a PP1/Glc7 phosphatase, which fails to maintain germline immortality at 25°C. GSP-2 is one of four PP1 catalytic subunits in *C. elegans* [22,23]. The PP1 phosphatase has roles in many cellular processes including mitosis, meiosis, apoptosis and protein synthesis [24].

Previously, GSP-2 has been shown to promote meiotic chromosome cohesion by restricting the activity of the Aurora B kinase ortholog AIR-2 to the short arms of *C. elegans* chromosomes during Meiosis I [25,26]. Here, we demonstrate that GSP-2 promotes germline immortality via a small RNA-mediated genome silencing pathway, implicating pairing and/or cohesion of meiotic chromosomes in genomic silencing.

## Results

### Identification of GSP-2 as a temperature-sensitive *mrt* mutant

In a screen for *mrt* mutants [2], one mutation that displayed a Temperature-sensitive defect in germ cell immortality, *yp14*, was tightly linked to an X chromosome segregation defect manifesting as a High Incidence of Males (Him) phenotype, such that 3.9% of *yp14* self-progeny were XO males, which significantly greater than the 0.05% male self-progeny observed in wildtype animals (Fig. 1A, p<.0001). The *yp14* mutation was mapped to Chromosome *III*, and whole genome sequencing revealed missense mutations in 6 genes within the *yp14* interval (Fig. S1A,B). Three-factor mapping of the *yp14* Him and Mrt phenotypes suggested that *yp14* might correspond to the missense mutation in *gsp-2* (Fig. 1C,D) or to a mutation in the G-protein coupled receptor gene *srb-11* (Fig. S1A,B).

**Figure 1:**
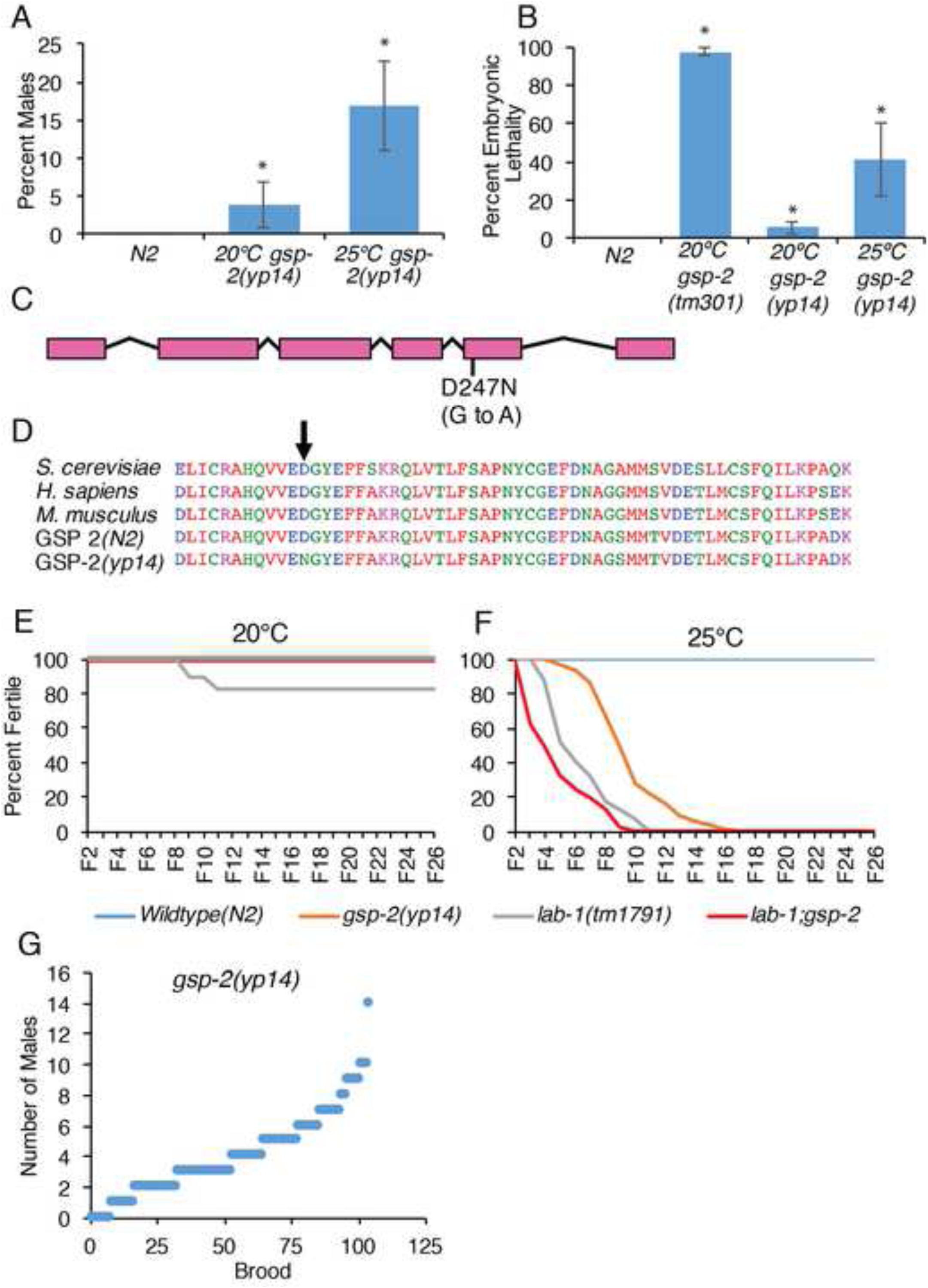
A hypomorphic mutation in *gsp-2* results in transgenerational sterility phenotype. **(A)** Incidence of males in *gsp-2(yp14)* was 3.9% at 20°C and increased to 16.8% at 25°C. **(B)** Progeny of *gsp-2(yp14)* animals grown at 20°C 25°C were 6% and 41% Embryonic Lethal, respectively, compared to 97% of *gsp-2(tm301)* progeny (N=20). **(C-D)** *gsp-2(yp14)* was identified to have a G to A mutation in exon 5 by whole genome sequencing. This results in a D to N amino acid substitution in a well conserved region of GSP-2. **(E)** When passaged at 20°C for many generations N2, *gsp-2(yp14), lab-1* and *lab-1; gsp-2(yp14)* did not exhibit a loss of transgenerational fertility. **(F)** *gsp-2(yp14)* and *lab-1* both exhibited loss of fertility at 25°C and were completely sterile by generation F17 and F11 respectively. A double mutant of *lab-1;gsp-2* went sterile slightly faster than the individual single mutants and were completely sterile by F10. (N≥40) **(G)** Analysis of incidence of males showed no jackpots of males at in *gsp-2(yp14)* animals. *P<.0001 by T-test.

To test whether the meiotic segregation phenotype of *yp14* was due to a mutation in *gsp-2*, we performed a non-complementation test with a deletion mutation in *gsp-2, tm301*. *yp14 / tm301* F1 heterozygous hermaphrodites gave rise to F2 male progeny at a frequency of 5.7% at 20°C, similar to the 3.8% male phenotype observed for *yp14* homozygotes (Fig. S1C). Thus, *tm301* failed to complement *gsp-2(yp14)* for its Him phenotype. In contrast, *gsp-2(tm301) / +* animals did not display a Him phenotype (Fig. S1C). *gsp-2(yp14)* strains also displayed 6% embryonic lethality at 20°C (Fig. 1B). Given that most forms of aneuploidy for the five *C. elegans* autosomes elicit embryonic lethality [27], the simplest explanation for the Him and Embryonic Lethal phenotypes of *gsp-2(yp14)* is that they are due to chromosome mis-segregation. Both of these phenotypes were exacerbated at 25°C (Fig. 1A,B), suggesting that *gsp-2(yp14)* has a chromosome segregation defect that may be mechanistically linked to its Mortal Germline phenotype (Fig. 1A,E). At both temperatures, the X chromosome non-disjunction defect was more pronounced than the embryonic lethality associated with non-disjunction of any of the five *C. elegans* autosomes (Table S1). *gsp-2(tm301)* null mutants immediately exhibited high levels of embryonic lethality at 20°C with a few F3 embryos that survive until adulthood (Fig. 1C), consistent with roles for PP1 in chromosome condensation and segregation during mitosis in several species [28–30]. Mutations that cause chromosome non-disjunction during mitosis occasionally lead to loss of an X chromosome during germ cell development, which could result in the stochastic appearance of XX hermaphrodites with high numbers of XO male progeny [27]. However, jackpots of XO males did not occur when *yp14* mutant hermaphrodites were isolated as single L4 larvae (Fig. 1G), implying that *yp14* is a separation-of-function mutation that specifically compromises the meiotic chromosome segregation function of GSP-2 without affecting its function in mitosis.

### GSP-2 promotes germline immortality at high temperature

GSP-2 is assumed to be localized to the long arms of meiotic chromosomes through binding to LAB-1, where it antagonizes AIR-2 (Aurora-B kinase) activity to regulate cohesion during meiotic prophase and Meiosis I [26,28,30]. In addition, LAB-1 is also present on mitotic chromosomes where it likely antagonizes AIR-2 activity [25]. In meiosis, LAB-1 fulfills the roles played by Shugoshin and Protein Phosphatase 2A in many other organisms, by protecting meiotic chromosome cohesion on the long arms in Meiosis I [25,31,32]. Once recruited by LAB-1, GSP-2 keeps REC-8, a meiosis-specific cohesin subunit, dephosphorylated to protect it from premature degradation and chromatid separation [25,26].

At 20°C, *gsp-2(yp14)* mutants remain fertile indefinitely, but at 25°C they exhibit sterility between generations F5 and F17 (Fig. 1E,F). We asked if a meiotic function of GSP-2 is relevant to germ cell immortality by first outcrossing a *lab-1* deletion with wildtype and re-isolating *lab-1* homozygotes in an effort to eliminate epigenetic defects that could have accumulated in the parental *lab-1* strain. Outcrossed *lab-1* mutants displayed a Mortal Germline phenotype at 25°C (Fig. 1E,F). We created *lab-1;gsp-2* double mutants, which remained fertile indefinitely when grown at 20°C but displayed a lightly accelerated number of generations to sterility at 25°C in comparison with *lab-1* mutants (Fig. 1E,F). Together, these results suggest that a meiotic function of GSP-2 that is directed by LAB-1 promotes germ cell immortality. Moreover, the weak Mortal Germline phenotype of *lab-1* single mutants at 20°C was suppressed by *gsp-2(yp14)* (Log Rank Test, p=.001).

### *gsp-2* and *lab-1* mutants display common germline defects at sterility

To investigate the cellular cause of transgenerational sterility in *gsp-2(yp14)* and *lab-1* mutants, we examined germline development in animals that became sterile after multiple generations. We previously reported that sterile *rsd-2* and *rsd-6* animals display a wide range of germline sizes, including many with few or no germ cells [12]. Interestingly, at the L4 larval stage, we found no significant difference between the germline profiles of *rsd-6* mutants at the generation of sterility and wild-type controls (Fig. 2H, Table S3). Most sterile generation L4 *gsp-2(yp14)* and *lab-1* mutant germlines were normal in size, though a small minority had a reduction in total germline length, resulting in a weak but significant difference in germline profile compared to wild-type (Fig. 2H, Table S3). Differentiating germ cell nuclei in spermatogenesis were observed for sterile generation L4 larvae for all strains (Fig. 2A,H). However, the germlines of two-day-old sterile *gsp-2, lab-1* and *rsd-6* mutant adults ranged in size from normal to a complete loss of germ cells (Fig. 2B-E,H), resulting in a strong significant difference when compared to wild-type controls (Table S3 p<1E-80). Given the similarities between the RNAi defective mutant *rsd-6* and *gsp-2*, we studied two additional small RNA silencing mutants. The germline profiles of adult sterile-generation *hrde-1* and *nrde-2* were similar to sterile *gsp-2(yp14)* animals (Fig. 2H, Table S2). Similar to *gsp-2* and *lab-1* mutants, *hrde-1* or *nrde-2* mutant L4 larvae that were poised to become sterile displayed predominantly normal-sized germlines (Fig. 2H). We previously showed that mutations in the cell death genes *ced-3* and *ced-4* partially rescued the empty and atrophy phenotypes observed for germlines of *rsd-2* and *rsd-6* small RNA silencing mutant adults [12], suggesting that apoptosis promotes germ cell atrophy as these animals develop from L4 larvae into adults.

**Figure 2:**
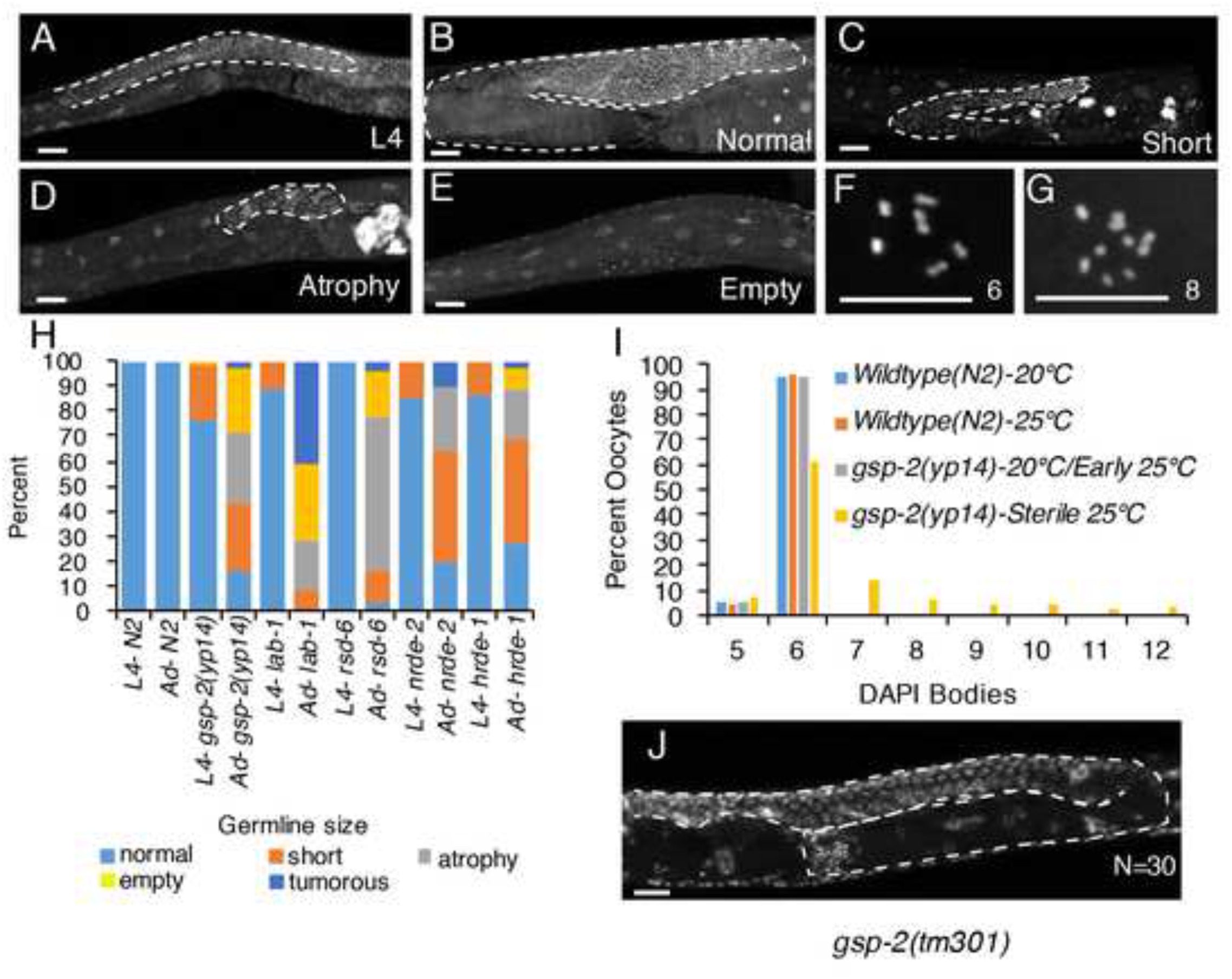
Germline defects occur in *gsp-2* and temperature-sensitive small RNA mutants at sterility. **(A-E)** Representative images of DAPI stained germlines passaged at 25°C. Germlines of either L4 (A) or adult control and sterile mutant animals were stained, and the germline size quantified as either normal (B), short (C), atrophied (D) or empty (E). **(H)** Germlines from *gsp-2(yp14), lab-1, rsd-6, nrde-2* and *hrde-1* mutants were examined and found have mostly normal morphology at the L4 stage but exhibited germline atrophy in adult animals (N≥98). **(F,G,I)** In addition to germline atrophy, *gsp-2(yp14)* animals displayed greater than the wildtype number of 6 DAPI bodies in oocytes at the generation at sterility in 32% of oocytes (N≥100). **(J)** 100% of adult *gsp-2(tm301)* animals displayed normal germline size by DAPI staining (N=30). P-values present in Table S2 and S3. Scale bar= 10um.

### Acute loss of GSP-2 does not cause small RNA mutant germline phenotypes

Previous work on GSP-2 in *C. elegans* reported that offspring of *gsp-2(tm301)* F2 homozygotes exhibited 98% embryonic lethality and that sterility occurred for F3 escapers [28,30]. To determine if acute loss of GSP-2 causes germline atrophy, we examined *gsp-2(tm301)* null mutants grown at 20°C and 25°C. We observed high levels of embryonic lethality for F3 embryos (97%), with a few escapers surviving to become uniformly sterile F3 adults that produced no F4 progeny [28] (Fig. 1B). These very high levels of embryonic lethality contrast with 41.6% embryonic lethality seen in *gsp2(yp14)* F8 animals grown at 25°C (Fig. 1B). *gsp2(tm301)* homozygous F2 animals and their few surviving F3 progeny showed normal germlines, with no morphological defects in germline size or development for either L4 larvae or young adults, which significantly differed from the germline profiles of *gsp-2(yp-14)* animals (Table S2, S3). Therefore, the late-generation sterility phenotype of *yp14* mutants is distinct from the fertility defects that occur in response to acute loss of GSP-2 in maternally depleted F3 deletion homozygotes.

### A meiotic recombination defect occurs at the generation of sterility

Mature *C. elegans* oocytes typically contain 6 bivalents (pairs of homologous chromosomes held together by crossovers), which can be scored as DAPI-stained bodies. Defects in meiosis can lead to the presence of univalents, which are observed as greater than 6 DAPI bodies per oocyte. We previously observed that small RNA nuclear silencing *mrt* mutants *rsd-2* and *rsd-6* displayed increased levels of univalents at sterility, which were not observed in either wildtype or in fertile *rsd-2* or *rsd-6* mutant late-generation animals grown at 25°C [12]. We measured the presence of univalents in *gsp-2(yp14)* and RNAi mutant strains and found that oocytes of control N2 wildtype or fertile *gsp-2(yp14)* worms grown at 20°C and 25°C almost always contained 6 DAPI bodies representing the 6 paired chromosomes (5 bodies are occasionally scored when bivalents that overlap spatially cannot be distinguished). However, when *gsp-2(yp14)* worms were passaged at 25°C until sterility, only 60% of oocytes containing 6 paired chromosomes with the other 40% containing 7 to 12 DAPI bodies (Fig. 2I). This increase in oocyte univalents was not present in fertile *gsp-2(yp14)* worms, even for late-generation 25°C strains close to sterility (Fig. 2I). We found no univalents in the null *gsp-2* allele *tm301*, either for F2 animals or for rare F3 escapers, consistent with previous observations [28,30]. Together these results emphasize that the germline phenotypes of sterile late-generation *gsp-2(yp14)* animals are distinct from those caused by acute zygotic or maternal loss of GSP-2 [25,26].

Given that LAB-1 and GSP-2 promote meiotic chromosome cohesion, we tested the hypothesis that dysfunction of other factors that promote meiotic chromosome cohesion might be sufficient to elicit germline atrophy. Sterile adults of mutant strains with defects in cohesion, *smc-3(t2553)* and in *coh-3(gk112); coh-4(tm1857)* double mutants [33–35] did not exhibit germline atrophy phenotypes observed in *gsp-2(yp14)* (Fig. S2, Table S2). Therefore, the late-generation sterility phenotypes of *gsp-2(yp14)* and small RNA mutants are not due to acute loss of meiotic chromosome cohesion.

### Increased expression of repetitive DNA in sterile *gsp-2(yp14)* and *rsd-6* mutants

One of the main functions of small RNA-mediated genomic silencing is to maintain silencing of repetitive elements and transposons in the germline, thereby protecting genomic integrity [15,19,36]. Previously, we reported that RNA expression of tandem repeat loci was upregulated in late-generation *rsd-2* and *rsd-6* mutants grown at 25°C [12]. Because similar levels of germline atrophy and univalents were observed in multiple small RNA pathway mutants and in *gsp-2(yp14)* at sterility, we asked if desilencing of tandem repeats occurred in *gsp-2(yp14)* mutants. We used RNA fluorescence *in situ* hybridization (FISH) to examine the expression of multiple repetitive elements. In wild-type controls grown at 25°C, we detected RNA from tandem repeat sequences using *CeRep59* sense and anti-sense probes in embryos but not in the adult germline or somatic cells, consistent with previous observations (Fig. 3A,B) [12].

**Figure 3:**
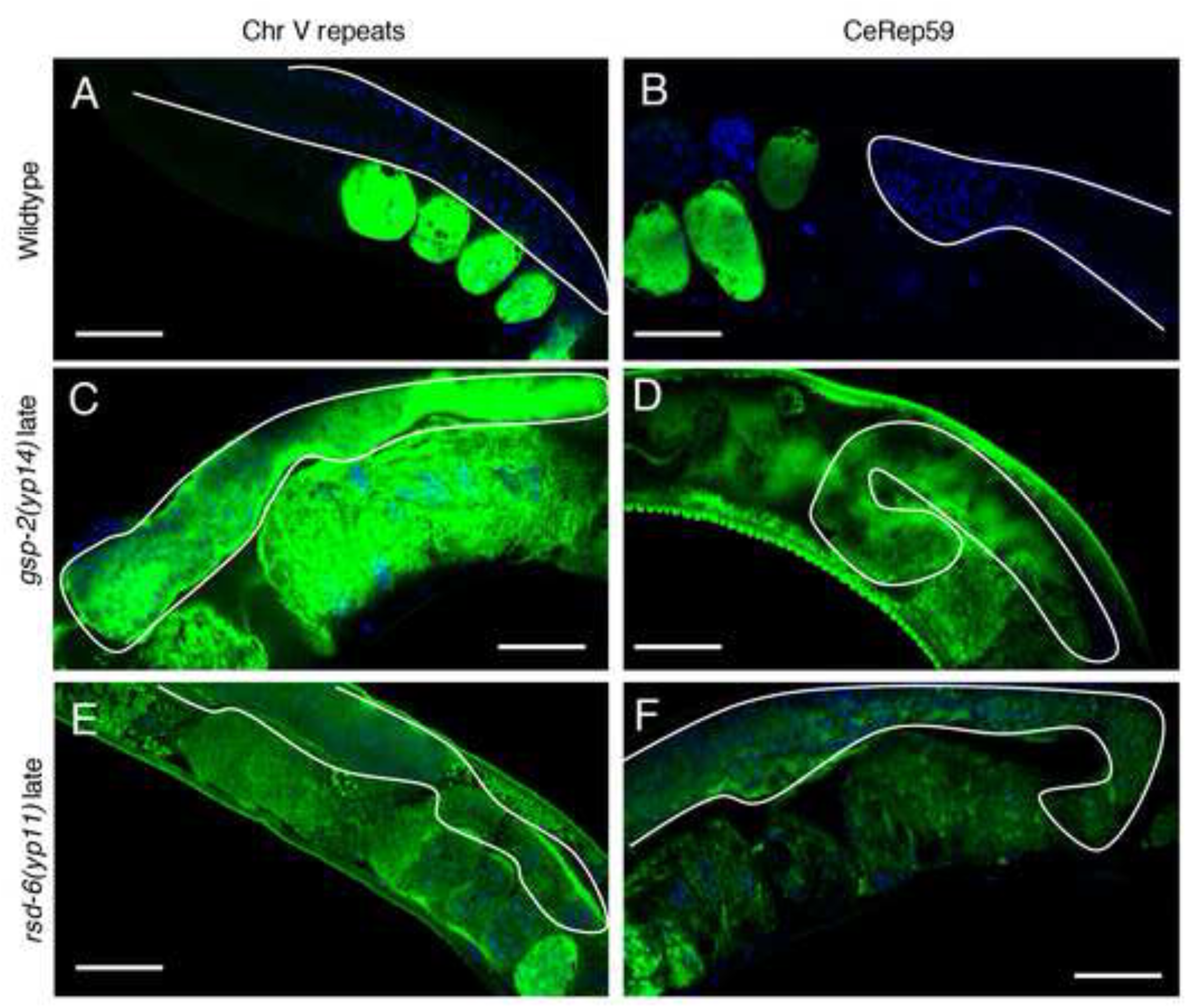
Repetitive regions in the genome are desilenced in *gsp-2(yp14)* animals. **(A-F)** Confocal images of Cy5-labeled RNA FISH probes (green) and DAPI-stained nuclei (blue). **(A,C,E)** RNA FISH probes show expression of Ch V repeats in the germlines of *gsp-2(yp14)* (C) and *rsd-6* (E) animals grown at 25°C and only embryonic expression in wildtype controls (A). **(B,D,F)** Probes against CeRep59 repeats reveal similar germline expression in *gsp-2(yp14)* (D) and *rsd-6* (F) animals and embryo-only expression in wildtype controls (B). All images were taken under the same condition. The germ line is outlined with white line. Scale bar= 30um

However, in late-generation *gsp-2(yp14)* and *rsd-6*, robust expression of tandem repeats was observed throughout the soma and germline of adult animals, indicating that tandem repeats become desilenced in these strains (Fig. 3C-F).

### Small RNA silencing components and *gsp-2* promote germ cell immortality

To study the relationship between *gsp-2(yp14)* and the small RNA nuclear silencing pathway, we created double mutants between *gsp-2(yp14)* and small RNA silencing mutants that display temperature-sensitive defects in germ cell immortality, *hrde-1, nrde-2* and *rsd-6.* Because *gsp-2(yp14)* is a hypomorphic allele, we predicted there would be a similar number of generations to sterility if it were functioning in the small RNA silencing pathway. For *gsp-2(yp14); hrde-1* and *rsd-6; gsp-2(yp14)*, we saw a modest decrease in the number of generations to sterility suggesting a slight additive effect (Fig. 4A,C, Log Rank test: p<.0001). In contrast, *nrde-2; gsp-2(yp14)* double mutants did not differ from the single mutants (Fig. 4B, Log Rank test: p=.06). Together, these results indicate that there is a weak additive effect on transgenerational lifespan when *gsp-2* is combined with *hrde-1* or *rsd-6*, but not when it is combined with *nrde-2*, which promotes the accumulation of stalled RNA polymerase II at loci that are targeted by small RNAs. The modest acceleration observed for some small RNA genomic silencing pathway and *gsp-2(yp14)* double mutants may be consistent with a recent report that indicated that epigenetic silencing marks typically overlap imperfectly for epigenomic silencing mutants, such that both shared and unique targets occur [15].

**Figure 4:**
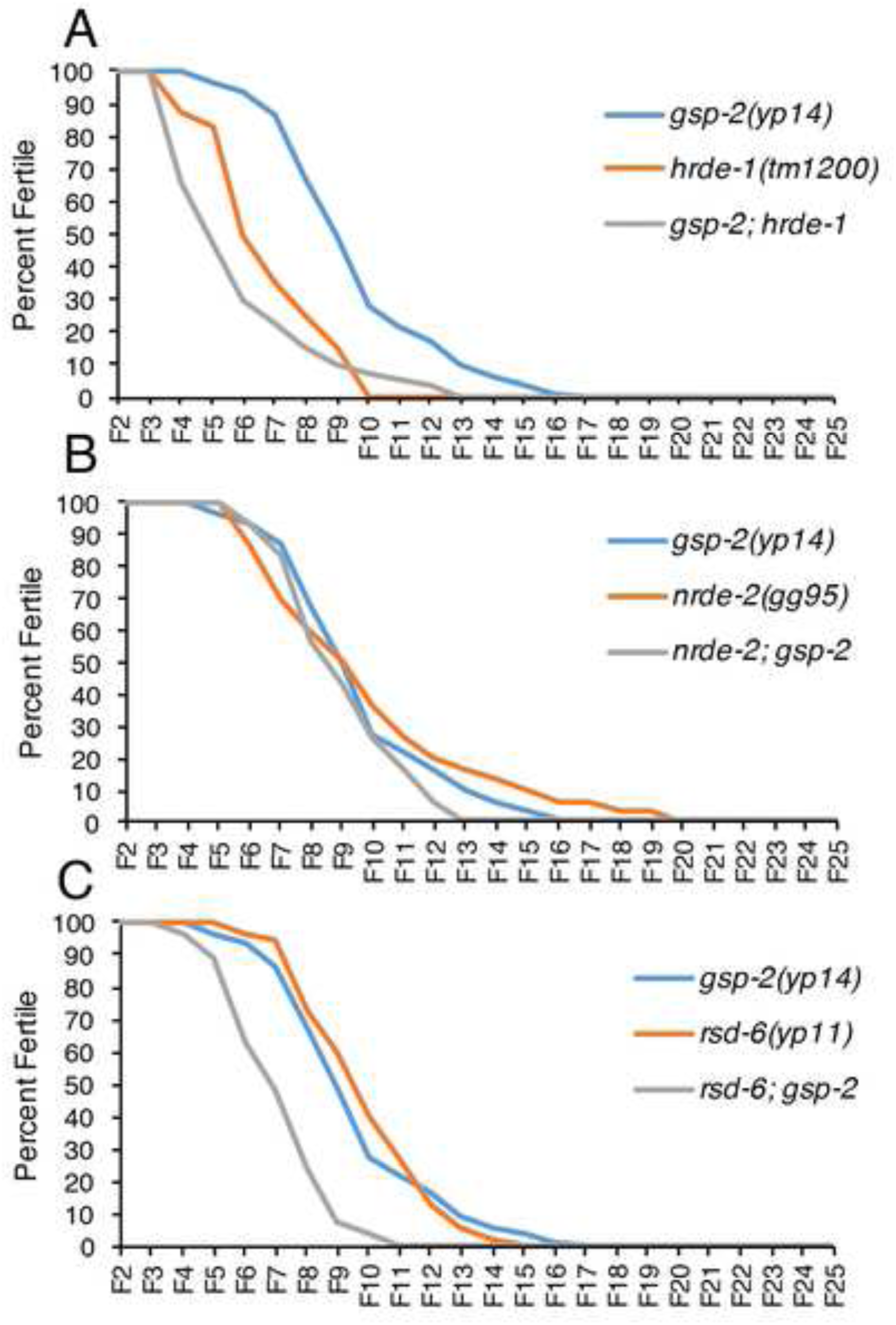
Temperature-sensitive small RNAi mutants exhibited similar times to sterility as *gsp-2(yp14)* at 25°C. Germline mortality assays all performed at 25°C **(A)** Both *gsp-2(yp14)* and *hrde-1* animals exhibit similar times to sterility while *gsp-2(yp14);hrde-1* double mutants display a slightly decreased time to sterility. p< .001**(B)** *gsp-2(yp14), nrde-2* and *rsd-6;gsp-2(yp14)* animals all go sterile in a similar number of generations. p=.06 **(C)** *gsp-2(yp14)* and *rsd-6* exhibit similar times to sterility while *gsp-2(yp14);rsd-6* double mutants become sterile at a slightly earlier generation. p<.001(N≥40). Significance was tested using a log rank test.

### *gsp-2(yp14)* and *lab-1* display increased histone H3 phosphorylation

Previously, increased H3S10 phosphorylation was observed when *gsp-2* is deficient [30]. The H3K9me and H3S10p marks can function as a phospho-methyl switch where H3S10 phosphorylation can block some epigenetic regulators, such as HP1, from accessing the adjacent H3K9me mark [37–39]. H3S10 phosphorylation was visible on the condensed chromosomes in the −1 to −3 oocytes in wildtype worms grown at 20°C and 25°C (Fig. 5A). In both early-and late-generation *gsp-2(yp14)* mutant oocytes, H3S10 phosphorylation increased when compared with wildtype controls, with increased levels on chromosomes (Fig. 5B,M). We observed increased levels of H3S10 phosphorylation in *lab-1* mutants (Fig. 5C,M), consistent with previous results [25].

**Figure 5:**
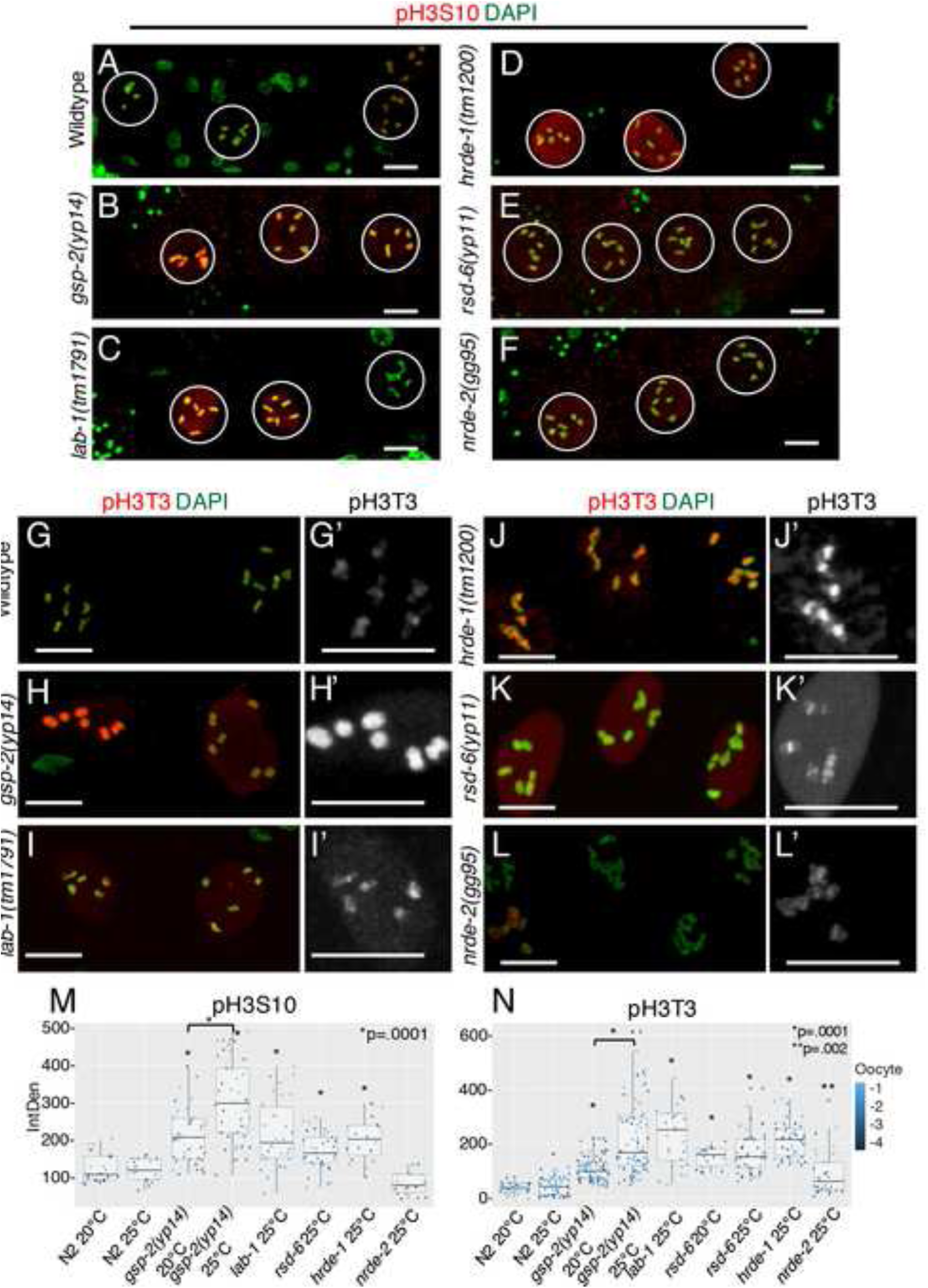
Increased histone phosphorylation is present in *gsp-2(yp14)* oocytes. **(A-F)** Day 2 late stage adults passaged at 25°C stained with an pH3S10 antibody (red) and DAPI marking the DNA (green). All samples were prepared at the same time and imaged using identical settings. **(A)** Wildtype control oocytes show low levels of H3S10p on condensed chromosomes. **(B)** *gsp-2(yp14)* oocytes have increased levels of H3S10p covering the entire chromosomes. **(C, F)** Both *lab-1* and *hrde-1* mutants also display increased levels of H3S10p but *nrde-2* **(D)** and *rsd-6* **(E)** do not. (N≥20**) (G-L)** Day 2 late generation or sterile adults passaged at 25°C stained with an pH3S10 antibody (red) and DAPI marking the DNA (green). All samples were prepared at the same time and imaged using identical settings. **(G)** Control oocytes show low levels of localized H3T3p on the short arms of the condensed chromosomes. **(H)** *gsp-2(yp14)* oocytes contain high levels of H3T3p that are expanded to cover the entire condensed chromosome. **(I-L)** *lab-1, hrde-1, rsd-6* and *nrde-2* all display varying levels of increased H3T3p staining compared to wildtype controls but localization to the short arms is still relatively normal. (N≥20) **(M)** Quantification of fluorescence intensity of H3S10p staining in N2, *gsp-2(yp14)* animals grown at 20°C and 25°C, *rsd-6, nrde-2, hrde-1*, and *lab-1* shows significant difference in staining intensity between N2 and mutants grown at the same temperature (except for *nrde-2*) and between *gsp-2(yp14)* mutants grown at 20°C and 25°C. (N≥20) **(N)** Quantification of fluorescence intensity of H3T3p staining in N2, *gsp-2(yp14)* animals grown at 20°C and 25°C, *rsd-6, nrde-2, hrde-1*, and *lab-1* shows significant difference in staining intensity between N2 and mutants grown at the same temperature and between *gsp-2(yp14)* mutants grown at 20°C and 25°C. (N≥20) Scale bar= 10um.

Increased levels of H3S10p also occurred in *lab-1, rsd-6*, and *hrde-1* but not in *nrde-2* mutants (Fig. 5C-F,M). Late-generation *gsp-2(yp14)* mutant animals grown at 25°C had a small but significant increase in H3S10 phosphorylation levels compared to *gsp-2(yp14)* mutant controls grown at 20°C (Fig. 5M). PP1 has been previously shown to dephosphorylate a number of histone amino acids, including H3T3 [40]. LAB-1 recruits GSP-2/PP1 to dephosphorylate H3T3, which represses the recruitment of Aurora B/AIR-2 to the long arms of *C. elegans* meiotic chromosomes, which separate in Meiosis II, thereby restricting Aurora B-mediated phosphorylation of H3S10 to meiotic short arms, which separate in Meiosis I [25,26,41]. We hypothesized that excess H3T3 phosphorylation could affect the ability of H3K4 demethylases to access H3K4 resulting in a Mrt phenotype similar to *rbr-2* or *spr-5* H3K4 demethylase mutants [42]. When we examined H3T3 phosphorylation in wildtype controls, staining was visible in the −1 to −3 oocytes and was localized to the short arms of meiotic chromosomes, the site of chromosome separation at Meiosis I (Fig. 5G-G’). The long arms, which do not exhibit staining, are analogous to the centromere in mono-centromeric organisms (Fig. 7). However, in *gsp-2(yp14)* mutants, H3T3p staining was expanded to cover the entire chromosome and was significantly brighter when images where taken under the same conditions (Fig. 5H-H’). In contrast to our H3S10p results, *lab-1* and the small RNA mutants *hrde-1, rsd-6* and *nrde-2* all exhibited increased H3T3 phosphorylation signal intensity in the −1 to −3 oocytes, although the signal was still localized properly to the short arms of the meiotic chromosomes (Fig. 5I-L,N).

Furthermore, there was a significant increase in H3T3 phosphorylation in the sterile generation of *gsp-2(yp14)* mutants compared to the earlier, fertile generation animals suggesting transgenerational accumulation of H3T3 phosphorylation (Fig. 5N).

Together, our results suggest that an increase in phosphorylation of H3T3 but not H3S10 consistently occurs in oocytes of *gsp-2* and small RNA silencing mutants. This defect is sensitive to temperature, as observed for the meiotic chromosome segregation and germ cell immortality defects of *gsp-2(yp14)* (Fig. 1E,F).

### Small RNA-mediated genome silencing is disrupted in *gsp-2(yp14)*

Multiple genes that regulate small RNA-mediated epigenomic silencing promote germ cell immortality, including three genes that are required for a form of transcriptional gene silencing termed nuclear RNA interference, *nrde-1, nrde-2* and *nrde-4* [10,12,43]. The response to a dsRNA trigger that targets *lin-26* is dependent on nuclear RNA interference [44]. Control wildtype and *gsp-2* mutant animals displayed a completely penetrant Embryonic Lethality phenotype in response to *lin-26* dsRNA, whereas nuclear RNAi defective mutant *nrde-2* and the RNAi defective mutant *rsd-6* did not (Fig. 6A), indicating that nuclear RNAi within a single generation is normal in the *gsp-2* mutant.

**Figure 6:**
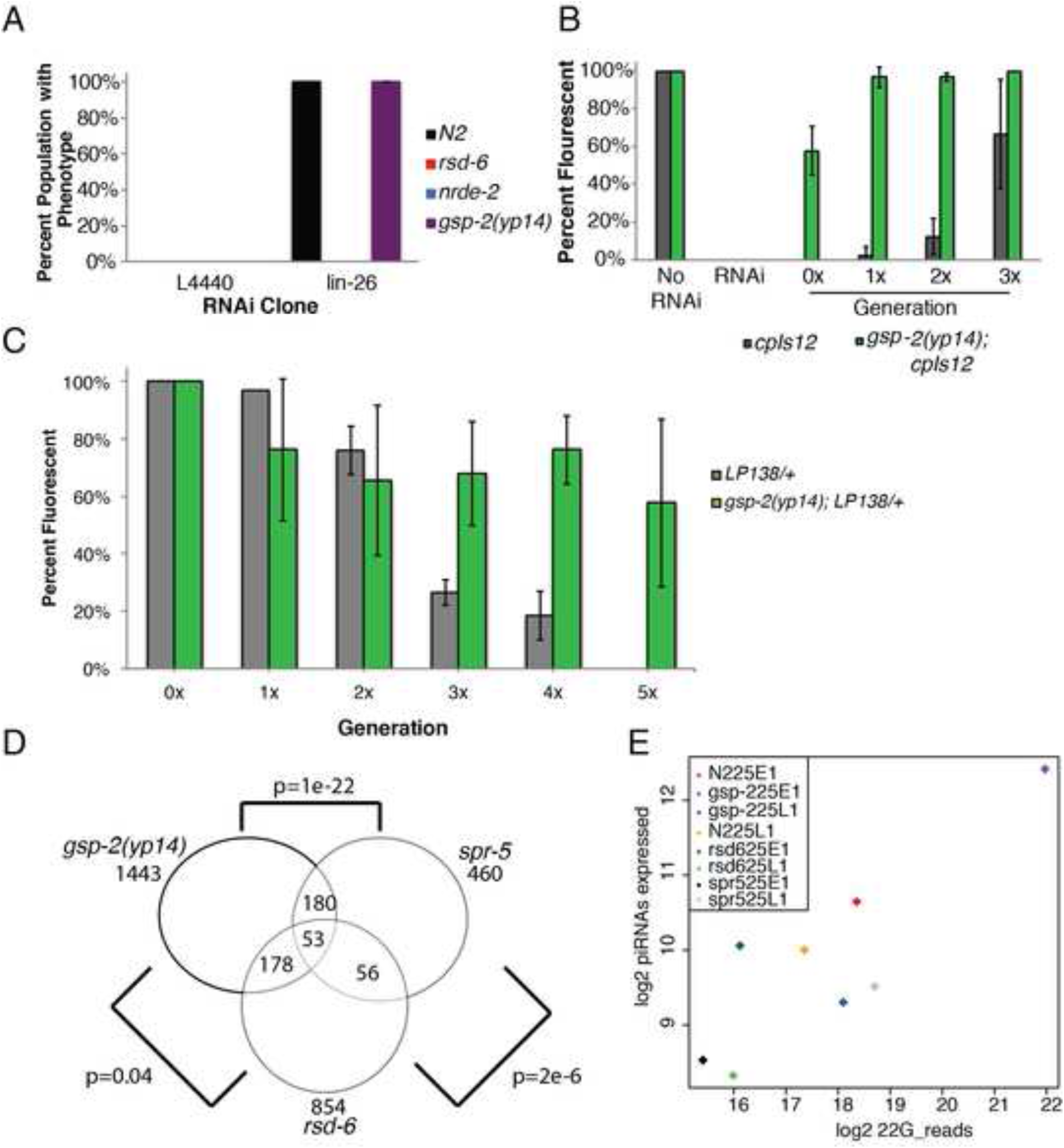
GSP-2 promotes multigenerational transgene silencing. **(A)** *gsp-2(yp14)* mutants do not exhibit single generation RNAi defects while *rsd-6* and *nrde-2* mutants are defective for single generation RNAi. **(B)** *cpIs12* remains undetectable for multiple generations after RNAi treatment. However, in *gsp-2(yp14);cpIs12* animals treated with RNAi cpIs12 only remains undetectable for one generation and by generation 3 exhibit close to wildtype levels of expression. **(C)** When LP138, a GFP transgene, is passaged as a heterozygote for multiple generations it is silenced in the germline. LP138 passaged as a heterozygote in a *gsp-2(yp14)* mutants results in only partial silencing over 5 generations suggesting defective heterozygous transgene silencing. **(D)** Comparison of small RNAs in *rsd-6, gsp-2* and *spr-5* mutants showing a great overlap in small RNA identity between *gsp-2* and *spr-5*. **(E)** Graph showing levels of piRNA expression in N2 controls, *rsd-6, gsp-2* and *spr-5* mutants at both early (E1) and late (L1) generations grown at 25°C.

**Figure 7:**
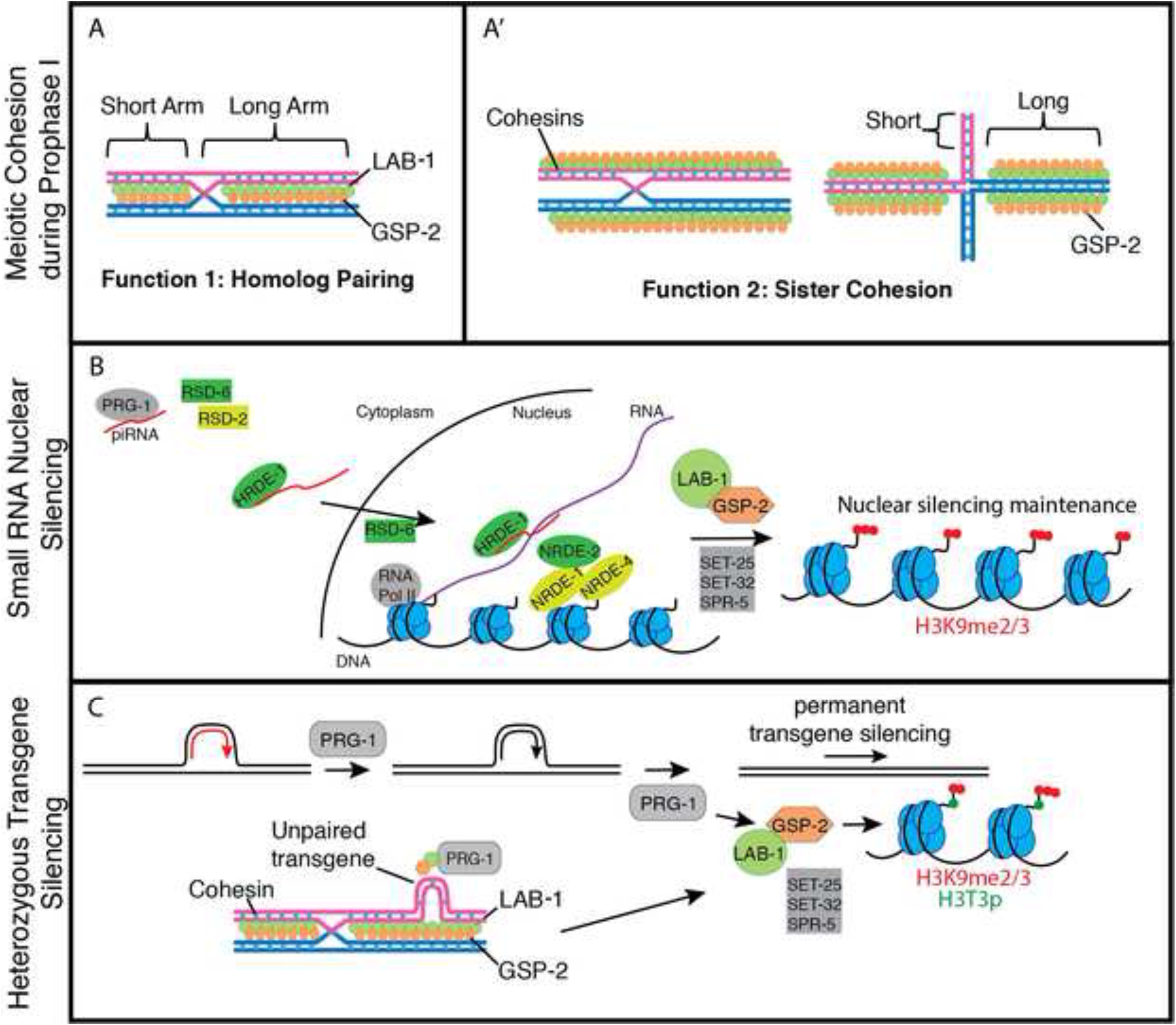
A model for the roles of GSP-2 and small RNA-mediated silencing in promoting germline immortality. We propose that both GSP-2 and small RNA-mediated silencing regulation the transgenerational inheritance of the epigenome. When these pathways are disrupted loss of epigenetic regulation can lead to germline atrophy. **A)** GSP-2/LAB-1 promote homolog pairing at the initiation of meiosis between homologous chromosomes during pachytene. **(A’)** GSP-2/LAB-1 promote sister chromatid cohesion during prophase. **B)** GSP-2 modulates small RNA silencing machinery promoting small RNA silencing potentially through histone dephosphoryation in a manner that promotes epigenetic silencing, **C)** Previous work has shown that PRG-1 is important for heterozygous transgene silencing (red=active transgene and black=silenced) in a similar manner to GSP-2. GSP-2/LAB-1 could function to silence heterozygous regions of DNA which disrupt meiotic pairing between homologs or cohesion between sister chromatids.

Small RNAs can trigger RNAi inheritance [10,13], where silencing of a gene in response to siRNAs can be transmitted for multiple generations. Transgenerational RNAi inheritance can occur when endogenous genes are targeted by dsRNA triggers [45], but this can also happen when GFP reporter transgenes are targeted by small RNAs derived from *GFP*. We tested the transgene *cpIs12* and found that it was silenced in response to GFP siRNAs and that silencing of this transgene was inherited for up to 4 generations after removal from the dsRNA trigger (Fig. 6B). In contrast, GFP expression in *gsp-2(yp14)*; *cpIs12* was initially silenced but silencing was not inherited over multiple generations (Fig. 6B), suggesting that *gsp-2* plays a role in RNAi inheritance.

Propagation of GFP or mCherry transgenes in the heterozygous state for multiple generations can elicit strong transgene silencing [21]. We found that the transgene *cpIs12* was progressively silenced in the population over the course of several generations until fully silenced by generation 5 (Fig. 6C). In contrast, when *cpIs12* was placed in a *gsp-2(yp14)* genetic background and propagated in a heterozygous state, we found that *cpIs12* was initially weakly silenced and that genomic silencing never became fully penetrant (Fig. 6C). Together, the above data indicate that *gsp-2* promotes transgenerational small RNA-induced genome silencing.

### Small RNA dysfunction in *gsp-2* mutants

Given that small RNA-mediated genome silencing is dysfunctional in *gsp-2(yp14)* mutants, we asked if small RNA populations were perturbed by preparing RNA from early-and late-generation wildtype, *gsp-2(yp14), rsd-6* and *spr-5* mutants grown at either 20°C or 25°C. *spr-5* encodes an H3K4 demethylase that promotes Germ Cell Immortality at 25°C, similar to *rsd-6* and *gsp-2* mutants [42]. Small RNA libraries were prepared and subjected to high throughput sequencing, and we then examined levels of 22G RNAs that are 22 nucleotides in length beginning with a 5’ guanine, as 22G RNAs are the major effectors of genomic silencing in *C. elegans* [5,46]. Analysis of the small RNA data revealed that *spr-5* and *rsd-6* share some genes with reduced levels of 22G RNAs with increasing generations, but there are other genes that show dissimilar behavior for each individual mutant. This suggests that *spr-5* may act both in conjunction with *rsd-6* and in a separate pathway to promote germline immortality. In contrast, 22G RNAs from *gsp-2(yp14)* showed strong similarities to those of *spr-5* mutants but showed little similarity to 22G RNA changes observed for *rsd-6* mutants (Fig. 6D and Fig. S3), suggesting that *gsp-2* may act in a separate pathway from *rsd-6*, in agreement with the results from epistasis experiments above. As a control, there is little coherent change in late-generation versus early generation N2 wildtype that overlaps with *gsp-2(yp14)* (Fig. S3). As germ cell immortality is promoted in part by primary siRNAs termed piRNAs that interact with the Piwi Argonaute protein PRG-1 [8], we also examined piRNA populations, which are enriched for 21 nucleotide RNAs that begin with a 5’ uracil (21U RNAs) [6,7,9] and found that these were normal (Fig. 6E).

## Discussion

We demonstrate for the first time that *gsp-2* and *lab-1* are required for germ cell immortality at 25°C as strains deficient for these proteins can be passaged for a number of generations before becoming sterile (Fig. 1C,D). Although PP1 is a general protein phosphatase with roles in a number of cellular processes including mitosis and meiosis [24], we identified a separation-of-function allele of *gsp-2* that displayed an X chromosome non-disjunction phenotype that was specific for meiosis (Fig. 1B,G). The incidence of both X chromosome loss and inviable embryos, which are likely aneuploid for autosomes, was exacerbated at high temperature (Fig. 1A,B), suggesting that this meiotic chromosome segregation defect could contribute to the temperature-sensitive nature of the Mrt phenotype of *gsp-2(yp14)*. Consistently, PP1/GSP-2 is recruited to meiotic chromosomes by the *C. elegans*-specific protein LAB-1, and we found that deficiency for *lab-1* elicited transgenerational sterility accompanied by adult germ cell degeneration phenotypes that were observed in sterile small RNA silencing mutants (Fig. 1F and Fig 2). Together, these results indicate that LAB-1 and GSP-2/PP1 define a step during meiosis that is critical for potentiating transgenerational small RNA-mediated genome silencing and germ cell immortality (Fig. 7A).

LAB-1 localizes to the interface between homologous chromosomes during pachytene, and LAB-1 recruits GSP-2 to nuclei during early stages of meiosis (Fig. 7A) (de Carvalho et al., 2008; Tzur et al., 2012). Moreover, deficiency for LAB-1 perturbs the pairing of homologous chromosomes during meiosis, which has been attributed to a defect in the cohesion of sister chromatids (Tzur et al., 2012). We suggest that a previously undescribed function of LAB-1/GSP-2, potentially related to the pairing of homologous chromosomes, may act at the interface between homologs to promote small RNA-mediated epigenomic silencing (Fig. 7A). We found that persistent transgenerational discontinuities in the pairing of homologous chromosomes during meiosis, caused by hemizygous transgenes, can promote transgene silencing in a manner that depends on GSP-2 (Fig. 6C, Fig. 7C). This hemizygous transgene silencing process occurs in a manner that depends *prg-1*/piRNAs as well as downstream factors that promote second siRNA biogenesis [21]. However, we found that piRNA levels were normal in *gsp-2(yp14)* mutants, and also that late-generation *gsp-2(yp14)* strains displayed changes in 22G RNA levels that were similar to those of *spr-5* histone H3K4 demethylase mutants but not to those of *rsd-6* small RNA biogenesis mutants (Fig. 6D and Fig. S3). Moreover, epistasis analysis indicated that there is a weak additive effect when *gsp-2* is combined with the nuclear Argonaute *hrde-1* or the small RNA biogenesis factor *rsd-6*, but no additive effect when *gsp-2* is combined with *nrde-2* (Fig. 4), which like *spr-5* functions downstream of siRNAs to promote transcriptional silencing [47]. The parallel with *spr-5* mutant small RNA levels suggests that GSP-2 help to integrate histone silencing modifications with the response to small RNAs (Fig. 7B). An intriguing possibility is that hemizygous transgenes could create a structural discontinuity between homologous chromosomes that alter the normal meiotic function of LAB-1/GSP-2, creating an environment where the chromosome silencing machinery can respond to small RNAs (Fig. 7C). Alternatively, the presence of a homologous allele could provide protection from silencing [48–50].

These possibilities are consistent with a role for small RNA-mediated genome silencing in responding to structures of *de novo* transposon insertions that present a threat to genomic integrity and that might create small hemizygous chromosome pairing aberrations during meiosis, which would be structurally analogous to hemizygous transgenes (Fig. 7C).

Consistent with our results, an allele of the *Drosophila* Protein Phosphatase 1 gene, *Su var (3) 6*, was identified as a suppressor of position-effect variegation, which relieves epigenetic silencing of a transcriptionally active gene that is placed adjacent to a segment of heterochromatin [29]. As position-effect variegation is promoted by small RNA-mediated genome silencing in plants and animals [51,52], we conclude that PP1 is likely to play a conserved role in this epigenomic silencing process. It has been suggested that the heterochromatin defect of *Su var (3) 6* mutants could reflect a direct role of PP1 in dephosphorylation of H3S10p, a mark that results in dissociation of Heterochromatin Protein 1 from heterochromatin [40,41]. Moreover, human PP1 has been shown to dephosphorylate H3T3p, this function is also carried out by *C. elegans* GSP-2 during meiosis [40,41]. One or both of these silencing marks could be relevant to meiotic small RNA-mediated genome silencing. Interestingly, the cohesin complex has been found to be recruited to silent chromatin, but in most cases has not been found to be necessary for genomic silencing [53]. As GSP-2 plays a role in protecting cohesin from being prematurely removed, it is likely localized to silent chromatin where it may play a role in maintaining silencing.

We observed increased H3T3 and H3S10 phosphorylation in mature oocytes of *gsp-2(yp14)* mutants as well as in small RNA genome silencing mutants. While the H3 phosphorylation defects that we observed were mildly sensitive to temperature, *gsp-2(yp14)* displayed more pronounced H3 phosphorylation defects that spread along the length of meiotic chromosomes, whereas small RNA mutants displayed proper H3 phosphorylation localization whose intensity was consistently stronger than for wildtype controls for H3T3 but not for H3S10 (Fig. 5). Given that LAB-1/GSP-1 localize to sister chromosomes in mature oocytes that are arrested in Meiosis I, where homologous chromosomes are held together only by crossovers, we propose that the role of LAB-1/GSP-2 in genome silencing may be at earlier stages of meiosis where the homologs are paired in a manner that might be capable of responding to discontinuities between homologs. We suggest that the changes in H3T3 phosphorylation of genome silencing mutants could reflect an indirect effect of dysregulation of heterochromatin, for example a response to altered levels of genome-wide levels of H3K9 methylation could affect the activity of protein that functions in the context of heterochromatin, such as the H3T3 kinase Haspin [43]. As H3T3 phosphorylation is constitutively disrupted in maturing oocytes of the genomic silencing mutants we studied, especially in *gsp-2* and *lab-1* mutants, it is also possible that this phosphorylation mark plays a role in genomic silencing itself.

While piRNAs and associated small RNA-mediated genome silencing have been well defined as functioning primarily in the germ line, our study suggests a novel connection between a meiotic process where Piwi-associated small RNAs scan homologous chromosomes for relatively small changes to their primary structure and germ cell immortality. This concept is well supported by work in other organisms, particularly in various fungi and *Drosophila*, where it has been shown that regions of heterozygosity are prone to silencing during meiosis in a small RNA dependent manner (reviewed by [20]). It is also possible that the GSP-2 protein phosphatase promotes small RNA-mediated genome silencing by directly modifying histones or a component of the genome silencing machinery that responds to small RNAs. In principle, our study defines a meiotic process that may intimately link transgenerational small RNA-mediated genome silencing with the structure of paired homologous chromosomes during meiosis. Given that endogenous small RNAs promote germ line stem cell maintenance, oogenesis and meiosis itself [54,55], we suggest that small RNA pathways and germ cell development have evolved to become mutually reinforcing processes.

## Materials and Methods

### Strains

All strains were cultured at 20°C or 25°C on Nematode Growth Medium (NGM) plates seeded with *E. coli* OP50. Strains used include Bristol N2 wild type, *gsp-2(tm301) III, gsp-2(yp14) III, lab-1(tm1791) I, cpIs12[Pmex-5::GFP::tbb-2 3’UTR + unc-119(+)] II, hrde-1(tm1200) III, rsd-6(yp11) I, nrde-2(gg95) II, rbr-2(tm1231) IV, smc-3(t2553) III, coh-4(tm1857) V, coh-3(gk112) V, air-2(or207) I, unc-32(e189) III, unc-13(e450) I, unc-24(e1172) IV.*

### Germline Mortality Assay

Worms were assessed for the Mortal Germline phenotype using the assay previously described [2]. L1 or L2 larvae were transferred for all assays. After passaging plates that yielded no additional L1 animals were marked as sterile. Log-rank analysis was used to determine differences of transgenerational lifespan between strains.

### DAPI staining

DAPI staining was performed as previously described. L4 larvae were selected from sibling plates and sterile adults were singled as late L4s, observed 24 hours later for confirmed sterility, and then stained 48 hours after collection.

### RNA FISH

DNA oligonucleotide probes coupled with a 5′ Cy5 fluorophore were used to detect repetitive element expression. The four probes used in this study were as follows: tttctgaaggcagtaattct, *CeRep59* on chromosome *I* (located at 4281435–4294595 nt); agaattactgccttcagaaa, antisense *CeRep59* on chromosome *I*; caactgaatccagctcctca, chromosome *V* tandem repeat (located at 8699155–8702766 nt); and gcttaagttcagcgggtaat, 26S rRNA. The strains used for RNA FISH experiments were *rsd-6(yp11), gsp-2(yp14)*, and N2 Bristol wild type. Staining was performed as described by Sakaguchi et al., 2014.

### Immunofluorescence

Adult hermaphrodites raised at 20°C or 25°C were dissected in M9 buffer and flash frozen on dry ice before fixation for 1 min in methanol at −20°C. After washing in PBS supplemented with 0.1% Tween-20 (PBST), primary antibody diluted in in PBST was used to immunostain overnight at 4 °C in a humid chamber. Primaries used were 1:500 pH3S10 (Millipore, 06570), 1:4000 pH3T3 (Cell Signaling, D5G1I, Rabbit) and 1:50 OIC1D4 (Developmental Studies Hybridoma Bank). Secondary antibody staining was performed by using an Cy3 donkey anti-mouse or Cy-5 donkey anti-rabbit overnight at 4°C. All images were obtained using a LSM 710 laser scanning confocal and were taken using same settings as control samples. Images processed using ImageJ. Intensity quantification was done by measuring total fluorescence in individual condensed chromosomes and subtracting the background levels obtained from mitotic nuclei as nucleoplasm levels varied greatly.

#### RNAi Assays

N2 wildtype, *gsp-2, rsd-6* and *nrde-2* animals were grown on *lin-26* RNAi clones and the progeny of 10 worms each were scored for Embryonic Lethality.

#### Transgene silencing assay

*cpIs12* and *gsp-2; cpIs12* worms were scored for GFP expression on NGM plates and then transferred to RNAi plates targeting GFP. The next generation (that was laid on GFPi plates) were scored for GFP expression and their sisters were removed and transferred back to NGM plates. Worms were propagated for multiple generations on NGM and scored each time for GFP expression. Both GFP reporter *gsp-2* doubles were created by marking with *dpy-17*.

#### Heterozygous transgene silencing

*cpIs12* was maintained as a heterozygote over *dpy-10 unc-4* while in *gsp-2; cpIs12* doubles, *gsp-2* remained mutant for the entire assay and *cpIs12* was balanced over *dpy-10 unc-4*.

### Genome sequence analysis

Paired sequence reads (2X100 nucleotide long) were mapped to the C. elegans reference genome version WS230 (www.wormbase.org) using the short-read aligner BWA [56]. The resulting alignment files were sorted and indexed with the help of the SAMtools toolbox [57]. The average sequencing depths for the mutant and wild-type N2 strains were 116x and 71x, respectively. Single-nucleotide variants (SNVs) were identified and filtered with SAMtools and annotated with a custom Perl script using gene information downloaded from WormBase WS230. Candidate SNVs in the mutant strain already present in the N2 strain were eliminated from further consideration.

The raw sequence data from this study have been submitted to the NCBI BioProject (http://www.ncbi.nlm.nih.gov/bioproject) under accession number PRJNA395732 and can be accessed from the Sequence Read Archive (SRA; https://www.ncbi.nlm.nih.gov/sra) with accession number SRP113543.

### Small RNA Sequence Analysis

5’ independent small RNA sequencing was performed as described previously [13], using one repeat for each time-point of N2 WT, *rsd-6* and *spr-5* at 25°C. Custom Perl scripts were used to select different small RNA species from the library. To map small RNA sequences to genes, reads were aligned to the *C. elegans* ce6 genome using Bowtie, Version 0.12.7, requiring perfect matches [58]. Data was normalized to the total number of aligned reads and 1 was added to the number of reads mapping to each gene to avoid division by zero errors. To map 22G sequences to transposons and tandem repeats, direct alignment to the transposon consensus sequences, downloaded from Repbase (Ver 17.05) or repeats obtained from the ce6 genome (WS190) annotations downloaded from UCSC as above, was performed using Bowtie allowing up to two mismatches and reporting only the best match. Uncollapsed fasta files were used for these alignments to compensate for the problem of multiple identical matches. Data was normalized to the total library size and 1 was added to the number of reads mapping to each feature to avoid division by zero errors. In order to analyze data from *rsd-2* mutants grown at 20°C [59], Fasta files were downloaded from the Gene Expression Omnibus and uncollapsed using a custom Perl script before aligning to transposons or tandem repeats as above. Analysis of data was carried out using the R statistical language [60].

## Acknowledgements

We thank Jonathan Hodgkin for mapping assistance and members of the S.A. laboratory for critical review of the manuscript. Some strains were provided by the Caenorhabditis Genetics Center, which is funded by National Institutes of Health Office of Research Infrastructure Programs (P40 OD010440). Work in the DM laboratory is funded by the Canadian Institute for Health Research. This work was supported by National Institute of Health grant numbers F32 GM120809 (K.B) and RO1 GM083048 (S.A). SA is a recipient of a Senior Scholar in Aging Award from the Ellison Medical Foundation. Work in PS laboratory is funded by the Medical Research Council. PS holds an Imperial College Research Fellowship.

---

**Supplemental Figure 1: Mapping and non-complementation test of *gsp-2(yp14)* (A)** Map of genomic region surrounding *gsp-2* on Chr. *III*. **(B)** Mapping of *gsp-2(yp14)* between *dpy-17* and *unc-32* on Chr. *III* placing *yp14* at -1.08. **(C)** Non-complementation test for Him phenotype between *gsp-2(yp14)* and *gsp-2(tm301)* showed an incidence of males of 5.7% at 20°C.

**Supplemental Figure 2: Loss of cohesion did not cause germline atrophy** DAPI staining and germline analysis showed no germline atrophy in *smc-3* and *coh-3; coh-4* mutants and minor defects in *air-2* animals suggesting loss of chromosome cohesion alone does not cause germline atrophy. (N=30)

**Supplemental Figure 3: *spr-5* and *gsp-2* show overlap in their small RNA populations (A)** Multigenerational inheritance assay using a second transgene *pkls32* in the background of *hrde-1* and *gsp-2* mutants. **(B-E)** Comparison of small RNAs in *rsd-6, gsp-2* and *spr-5* mutants: **(B)** *rsd-6* vs *gsp-2*, **(C)** *spr-5* vs *gsp-2*, **(D)** *rsd-6* vs *spr-5* and **(E)** N2 vs *gsp-2*.

**Supplemental Table 1.** Expected vs Observed Embryonic Lethality.

|  | Males | Exp Dead Emb | Exp Live Worms | Obs Dead Emb | Obs Live Worms | p-value |
| --- | --- | --- | --- | --- | --- | --- |
| **20°C *gsp-2(yp14)*** | 164 | 820 | 5290 | 88 | 1492 | <0.0001 |
| **25°C *gsp-2(yp14)*** | 155 | 775 | 964 | 481 | 1380 | <0.0001 |

**Supplemental Table 2:** P-values for adult germline defects in *gsp-2* and temperature-sensitive small RNA mutants.

|  | *N2* | *gsp-2(yp14)* | *lab-1* | *gsp-2(tm301)* | *rsd-6* | *nrde-2* | *hrde-1* | *smc-3* | *coh-3/4* | *air-2* |
| --- | --- | --- | --- | --- | --- | --- | --- | --- | --- | --- |
| *N2* | 1 | 1.008E-85 | 1.5E-90 | 1 | 2E-85 | 1.3E-67 | 2E-59 | 1 | 1 | 8.1E-55 |
| *gsp-2(yp14)* | 1E-85 | 1 | 5.1E-17 | 1.646E-45 | 8E-06 | 1.4E-06 | 0.0154 | 1.2E-40 | 6.7E-41 | 5.4E-23 |
| *lab-1* | 2E-90 | 5.13E-17 | 1 | 2.2041E-47 | 5E-11 | 1.2E-16 | 8E-17 | 7.3E-43 | 4.4E-43 | 1.8E-19 |
| *gsp-2(tm301)* | 1 | 1.646E-45 | 2.2E-47 | 1 | 1E-43 | 8.2E-32 | 9E-27 | 1 | 1 | 8.2E-25 |
| *rsd-6* | 2E-85 | 7.718E-06 | 4.9E-11 | 1.3493E-43 | 1 | 3.4E-12 | 9E-11 | 4E-39 | 2.5E-39 | 3.5E-29 |
| *nrde-2* | 1E-67 | 1.383E-06 | 1.2E-16 | 8.175E-32 | 3E-12 | 1 | 0.248 | 7.5E-28 | 4.9E-28 | 8.4E-08 |
| *hrde-1* | 2E-59 | 0.01536 | 7.8E-17 | 9.4732E-27 | 9E-11 | 0.248 | 1 | 4.1E-23 | 2.7E-23 | 7.6E-10 |
| *smc-3* | 1 | 1.151E-40 | 7.3E-43 | 1 | 4E-39 | 7.5E-28 | 4E-23 | 1 | 1 | 1.9E-21 |
| *coh-3/4* | 1 | 6.658E-41 | 4.4E-43 | 1 | 2E-39 | 4.9E-28 | 3E-23 | 1 | 1 | 1.3E-21 |
| *air-2* | 8E-55 | 5.438E-23 | 1.8E-19 | 8.1847E-25 | 4E-29 | 8.4E-08 | 8E-10 | 1.9E-21 | 1.3E-21 | 1 |

**Supplemental Table 3:**
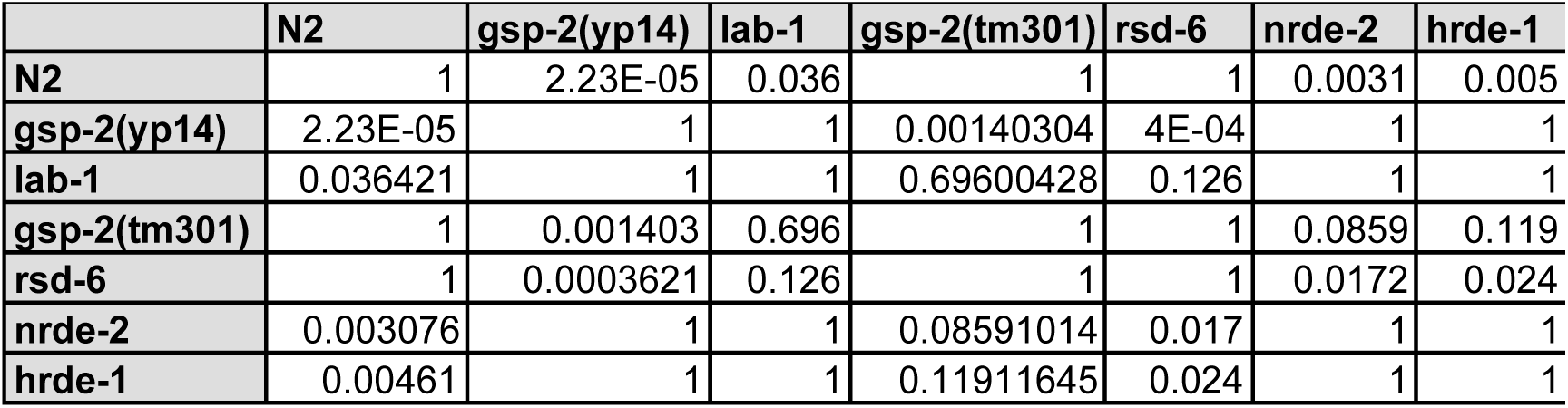
P-values for L4 germline defects in *gsp-2* and temperature-sensitive small RNA mutants.

